# Introduction of the *Aspergillus fumigatus* α-1,2-mannosidase MsdS into *Trichoderma reesei* leads to abnormal polarity and improves the ligno-cellulose degradation

**DOI:** 10.1101/2020.04.07.031062

**Authors:** Prakriti Sharma Ghimire, Haomiao Ouyang, Guangya Zhao, Mingming Xie, Hui Zhou, Jinghua Yang, Cheng Jin

**Affiliations:** State Key Laboratory of Mycology, Institute of Microbiology, Chinese Academy of Sciences, Beijing 100101, China; University of Chinese Academy of Sciences, Beijing 100101, China; National Engineering Research Center for Non-food Bio-refinery, Guangxi Academy of Sciences, Nanning 530007, Guangxi, China; Himalayan Environment Research Institute (HERI), Bouddha-6, Kathmandu, Nepal

**Keywords:** α-mannosidase, N-glycans processing, *Trichoderma reesei*, cellulase, protein secretion

## Abstract

α-1,2-Mannosidase is an important enzyme essential for N-glycan processing and plays a significant role in the biosynthesis and organization of fungal cell wall. Lacking of α-1,2-mannosidase leads to cell wall defect in yeast and filamentous fungi. *Trichoderma reesei* is known to be non-toxic to human, and its N-glycan on secreted glycoprotein is Man_8_GlcNAc_2_. To evaluate the significance of the N-glycan processing in *T. reesei*, in this study *Aspergillus fumigatus* α-1, 2-mannosidase MsdS, an enzyme that cleaves N-linked Man_8_GlcNAc_2_ in Golgi to produce Man_6_GlcNAc_2_ on secreted glycoprotein, was introduced into *T. reesei*. The *msdS*-expressing strain Tr-MsdS produced a major glycoform of Man_6_GlcNAc_2_ on its secreted glycoproteins, instead of Man_8_GlcNAc_2_ in the parent strain. Although the cell wall content of *msdS*-expressing strain Tr-MsdS was changed, it appeared that the cell wall integrity was not affected. However, phenotypes such as increased conidiation, multiple budding and random branching were observed in strain Tr-MsdS. In addition, expression of MsdS into *T. ressei* also affected protein secretion and improved the ligno-cellulose degradation of *T. reesei*. Our results indicate that processing of the N-glycan is species-specific and plays an important role in protein secretion in *T. reesei*, specially cellulases. Also, our results provide a new strategy to improve cellulases production by interfering the N-glycan processing in *T. reesei*.

**Importance:** For the first time, the N-glycan processing is shown to play an important role in polarized growth and protein secretion in *T. reesei*. In addition, our results show that alterated N-glycan processing enhances cellulose degradation, which provides a strategy to improve cellulases production in *T. reesei*.

## Introduction

*Trichoderma reesei* (syn. *Hypocrea jecorina*), a mesophilic filamentous fungus identified as GRAS (generally recognized as safe) status by FDA (U.S. Food and Drug Administration) (Kuhls et al. 1996; Schuster and Schmoll, 2010), is able to produce a wide range of extracellular enzymes in large quantities. *T. reesei* possesses protein post-translational modification and grows faster than plant, insect or mammalian cells, therefore it has been thought as an attractive expression host. However, the expression of heterologous protein is less efficient than the expression of endogenous proteins in *T. reesei* and, therefore, efforts have been made to improve the production and secretion of heterologous proteins (Kredics et al, 2014; Kiiskinen et al, 2004; Eades and Hintz, 2000).

Enzymes secreted by *T. reesei* are glycosylated by both *N*-glycan and O-glycan (Beckham et al, 2010). It has been shown that O-glycosylation affects not only the function of these enzymes, such as proteolytic resistance, thermostability and cellulose binding (Chen et al, 2014), but also their expression and secretion (Gorka-Niec et al, 2008; Gorka-Niec et al, 2011). Interestingly, either increase or decrease of O-glycosylation level leads to an increase of secreted proteins (Agaphonov et al, 2005; Perlinska-Lenart et al, 2006). However, it keeps unknown how N-glycan processing affects protein secretion.

In eukaryotic cells, secreted proteins are synthesized in the endoplasmic reticulum (ER) and transported to Golgi apparatus. In the course of the secretory pathway, N-glycosylation is initiated in the lumen of the ER by transferring of Glc_3_Man_9_GlcNAc_2_ from dolichol pyrophosphate to the nascent polypeptides. Once the Glc_3_Man_9_GlcNAc_2_ is transferred to proteins, N-Glycan processing is initiated sequentially in the ER and Golgi by two ER α-glucosidases and various 1,2-α-mannosidases (Kornfeld and Kornfeld, 1985; Lehle et al, 2006). In mammalian cells, Man_9_GlcNAc_2_ is converted to Man_5_GlcNAc_2_ by the action of ER and Golgi α- mannosidases, and Man_5_GlcNAc_2_ is the precursor for complex, hybrid, and high-mannose type of N-glycans (Kornfeld and Kornfeld, 1985). Golgi α-mannosidase is a Class-I α-mannosidase responsible for cleavage of α-1,2-linked D-mannoses (Van Petegem et al, 2001; Maras et al, 2000; Moremen et al, 1994). In vitro analysis reveals that this enzyme converts Man_8_GlcNAc_2_ or a mixture of Man_6-9_GlcNAc_2_ oligosaccharides to the respective Man_5_GlcNAc_2_ structures (Maras et al, 2000; Li et al, 2008).

Previously we have shown that in filamentous fungi *Aspergillus fumigatus* -mannosidase MsdS is responsible for processing of N-linked Man_8_-_9_GlcNAc_2_ to Man_6_GlcNAc_2_, which is the glycoform on secreted glycoproteins (Li et al, 2008). Deletion of the α-mannosidase gene *msdS* results in a conversion of N-glycan attached to secreted glycoproteins from Man_6_GlcNAc_2_ to Man_8_GlcNAc_2_ and causes defective cell wall and abnormal polarity (Li et al, 2008). On the other hand, it is interesting to note that the N-glycan on secreted glycoproteins of *T. reesei* is Man_8_GlcNAc_2_ (García, R. et al, 2001), which is the same as that of the *A. fumigatus msdS*-knockout mutant, suggesting that N-glycan processing is different in these two species.

In attempt to evaluate the effect of the altered N-glycan processing on *T. reesei*, in this study a *msdS*-expressing strain Tr-MsdS was constructed by introducing of the *A. fumigatus msdS* gene into *T. reesei*. As expected, the main N-glycan glycoform was converted from Man_8_GlcNAc_2_ to Man_6_GlcNAc_2_ in strain Tr-MsdS. Analysis of Tr-MsdS strain revealed that expression of MsdS led to abnormal polarity and altered protein secretion.

## Results

### Introduction of the *A. fumigatus msdS* gene into *T. reesei*

The *A. fumigatus msdS* gene (α-1,2-mannosidase) consist 1512 nucleotides and encodes a polypeptide of 503 amino acids with molecular mass of 55.44 kDa. The conserved domain of MsdS protein contains one functional domain that belongs to Glycosyl hydrolase family 47, a residues of the N-linked Man_9_GlcNAc_2_. The blast score of MsdS exhibits 82.0% amino acid identity with *A. clavatus α*-mannosidase, 77.8 % with *A. oryzae α*-mannosidase, 73.7 % with *A. niger*, and 48.9% with *H. jecorina*.

As described under Materials and Methods, an expression vector (pGM) containing *gpda* promoter and *msdS* gene was constructed (Fig.1). By transforming of the expression vector pGM into *T. reesei* TU-6 strain, a *msdS*-expressing strain Tr-MsdS was obtained. RT-PCR analysis revealed that the *msdS* gene was expressed in Tr-MsdS strain but not its parent strain (P<0.001) (Fig.2A). To detect if MsdS protein was expressed in Tr-MsdS strain, we prepared anti-MsdS antibody using recombinant MsdS expressed in *E. coli* (Li et al, 2008), however, the specificity of the antibody was not good (data not shown). Therefore, we further analyzed proteins extracted from Tr-MsdS strain with LC-MS/MS. To this end, the secreted proteins of Tr-MsdS strain were separated by polyacrylamide gels and protein bands corresponding to 40-60 kDa were subjected to LC-MS/MS analysis. As a result, 16 peptides were identified as fragments of MsdS (gi159131578) (Fig.S1 and Fig.S2). Although a MsdS homolog (TRIREDRAFT_45717) was found in *T. reesei* (Fig.S3), no peptide was identified. These results clearly demonstrated that MsdS was successfully expressed in Tr-MsdS strain, while the *T. reesei* MsdS homolog was not expressed.

**Fig.1.**
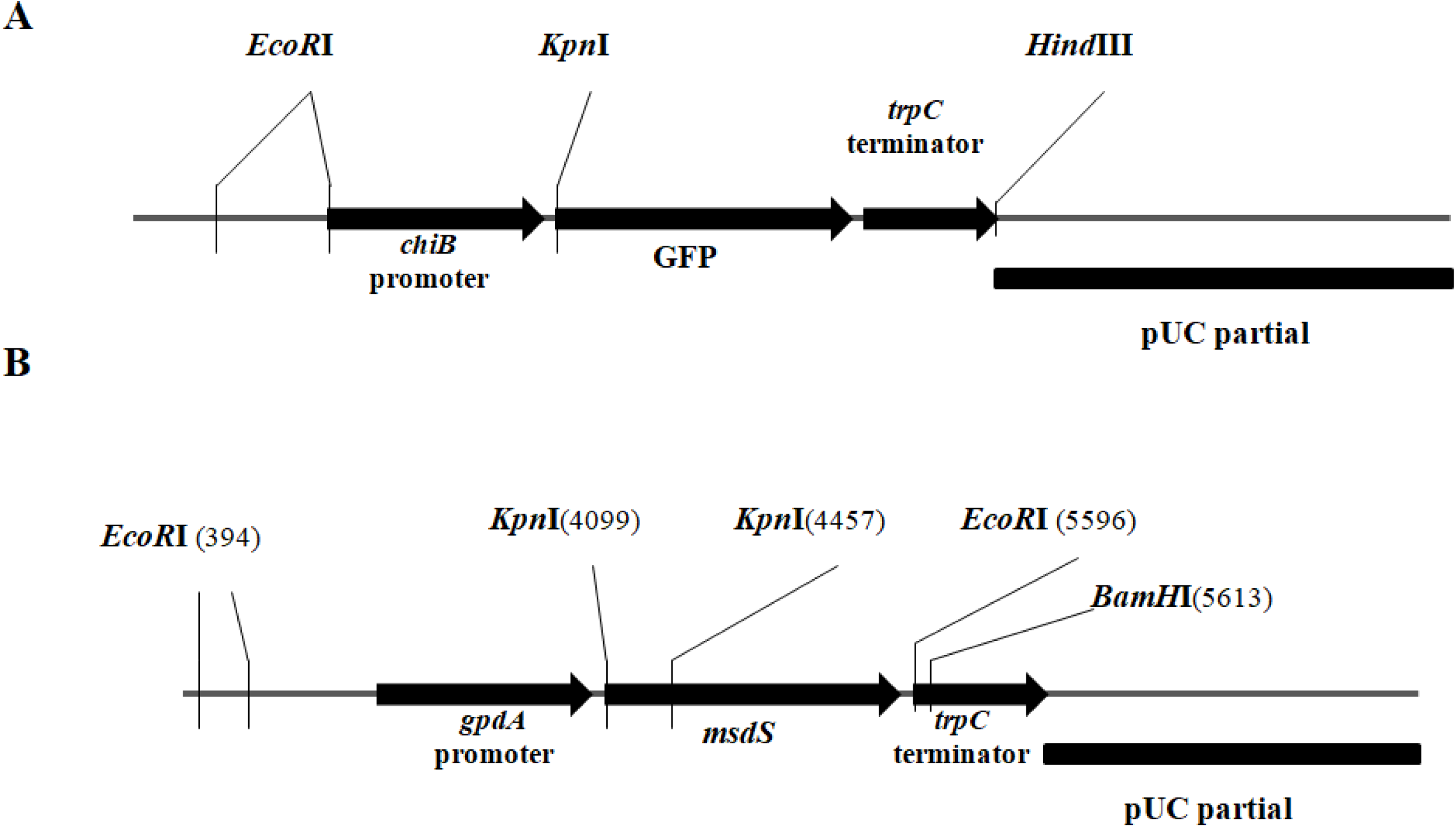
Schmatic of the *msdS*-expressing vector construction. A, a parent vector containing *chiB* promoter, GFP fragment and terminator *trpC*. B, a new recombinant vector by replacing *chiB* promoter with *gpdA* promoter and GFP fragment with *msdS* gene. The insertion of *gpdA* promoter was done at restriction site *EcoR*I and *Kpn*I, while insertion of the *msdS* gene was done at *Kpn*I and *BamH*I restriction site.

**Fig.2.**
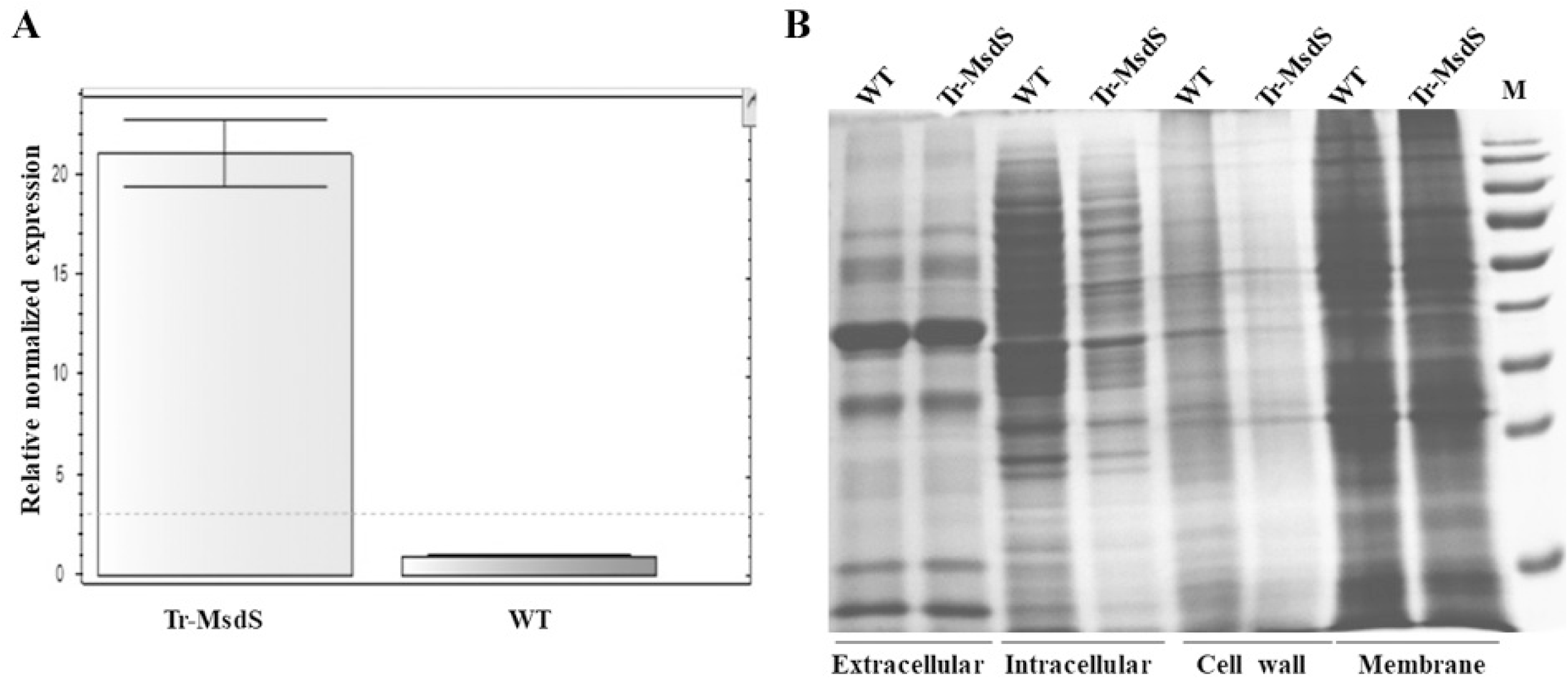
Expression of the *msdS* gene in *T. reesei*. In A, expression of the *msdS* gene was measured with total RNA from the wild-type (WT) or Tr-MsdS cultured at 32°C. RNA extraction and cDNA synthesis were carried out as described in method section. Results are repeated three times and presented as mean ± SD. In B, extracellular, intracellular, membrane and cell wall protein were was extracted as mentioned in method and separated on SDS-PAGE.

### Phenotype of Tr-MsdS strain

The growth kinetics of *T. reesei* strains were studied on solid media by measuring colony diameter. The result showed that 28-32°C were the favorable temperatures for both parent and Tr-MsdS strains, however Tr-MsdS strain was observed to be slightly temperature-sensitive at 42°C as compared with its parent strain (Fig.S4). Additionally, at 32 °C, the conidia produced by parent strain was more than that produced by Tr-MsdS strain (47%, 61% and 38% at 12 h, 24 h and 48 h, respectively), whereas after 72 h the conidia produced by Tr-MsdS strain appeared higher. This suggests that expression of MsdS leads to a slightly slower growth rate at early phase (Table 1).

**Table 1.** Conidia produced by Tr-MsdS strain at 32°C.

| Time (h) | Strain | Spore count (*10<sup>4</sup>) | Time (h) | Strain | Spore count (*10<sup>4</sup>) |
| --- | --- | --- | --- | --- | --- |
| 12 | WT | 3.5±1.3 | 96 | WT | 402.3±16.6 |
|  | Tr-MsdS | 1.3±0.5 |  | Tr-MsdS | 453.8±11.7 |
| 24 | WT | 10.3±1.7 | 120 | WT | 531.8±28.7 |
|  | Tr-MsdS | 2.5±1 |  | Tr-MsdS | 552.8±10.4 |
| 48 | WT | 16±2.5 | 144 | WT | 658±28.4 |
|  | Tr-MsdS | 7.3±2.1 |  | Tr-MsdS | 897.5±35.5 |
| 72 | WT | 267.3±24.8 | 168 | WT | 644.5±41.3 |
|  | Tr-MsdS | 376±12.0 |  | Tr-MsdS | 891.8±29.4 |

Same amount of conidia was spotted on solid complete medium and incubated at 30°C. Conidia were harvested at the specified times, resuspended in ddH<sub>2</sub>O, and counted under microscopy using haemocytometer. A triplet of each strain was counted. The same experiment was repeated twice.

As shown in Fig.3A, in the presence of antifungal reagent, hyphal growth of Tr-MsdS strain was not affected at 28°C, 32°C or 37°C. When observed under transmission electron microscope (TEM) (Fig.3B), Tr-MsdS strain grown at 32°C showed 30% more thickened hyphal cell wall as compared with its parent strain. When the temperature was raised to 37°C, the thickness of the Tr-MsdS mycelia cell wall was only 13 % more as compared with its parent strain. In addition, the conidial cell wall of Tr-MsdS strain formed at 32°C was 9% thicker than at 37°C, whereas the cell wall of parent strain showed 27% less thick at 37°C as compared with 32°C. Interestingly, Tr-MsdS strain showed less dense and irregularly scattered filamentous material around spore and hypha, whereas its parent strain showed denser, regular and prominent filamentous material surrounding spore and hyphae. These results indicated that hyphal cell wall thickness was changed in Tr-MsdS strain and altered at 37°C.

**Fig.3.**
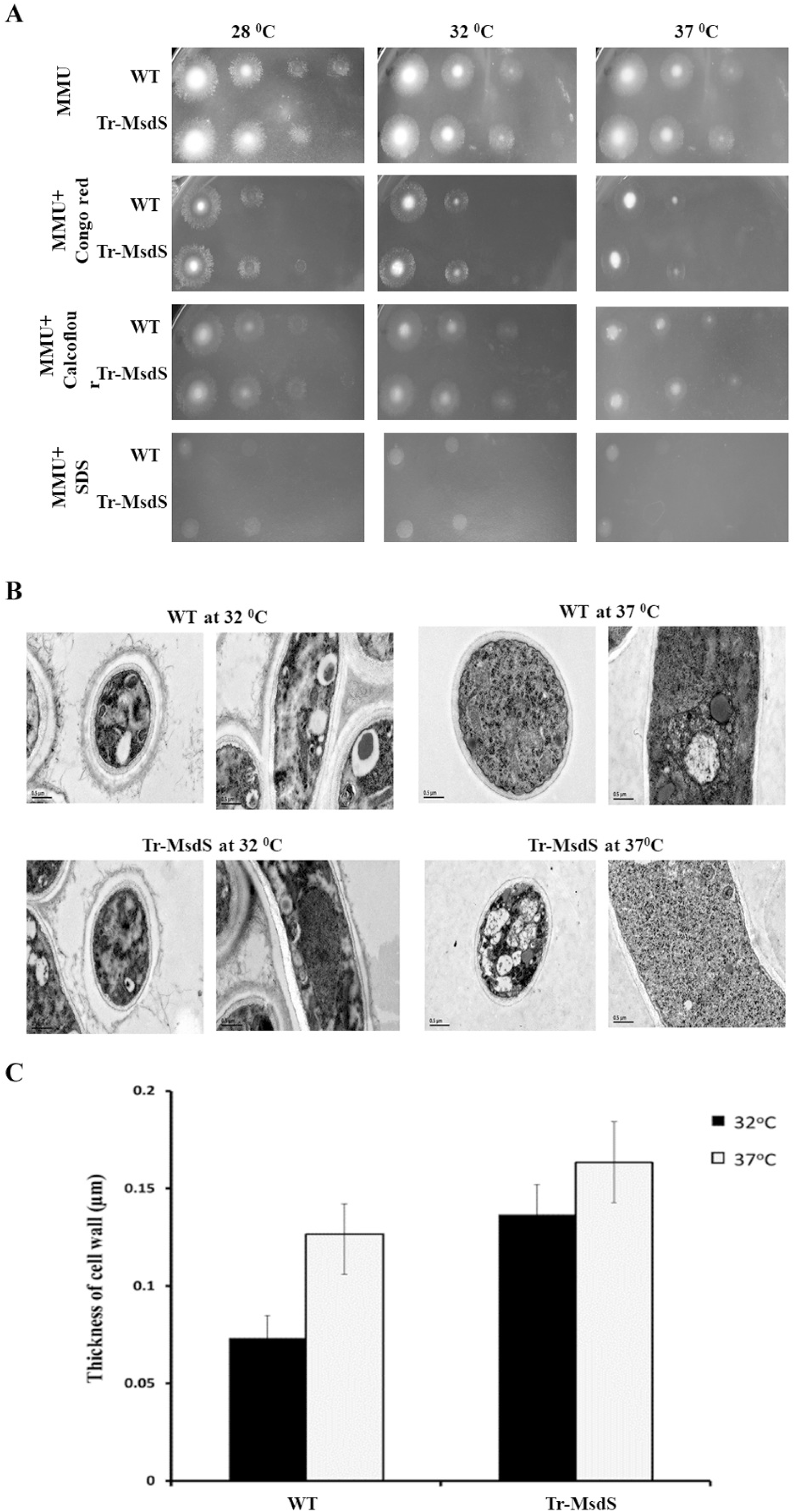
Phenotype and morphology of Tr-MsdS strain. In A, sensitivity to cell wall perturbing compounds was carried out by spotting conidiophores on solid MMU plates supplemented with 100 µg/ml calcofluor white, 150 µg Congo red or 40 µg SDS and incubating at 28°C, 32°C and 37°C. In B, the mycelia and conidial cells were fixed as described in method section and examined with transmission electron microscopy (TEM) (JEM-1400). In C, thickness (µm) of cell wall was measured under TEM.

The cell wall components including glycoprotein, glucan and chitin were further analyzed (Table 2). The glycoprotein content was reduced by 30-40% in Tr-MsdS strain at both 32°C and 37°C. The cell wall chitin in Tr-MsdS strain was reduced by 15% at 32°C and by 10% at 37°C. Interestingly, α/β-glucan was found to be increased by 25% in Tr-MsdS strain. These phenomena suggested that the expression of MsdS resulted in an increase of α/β-glucan in *T. reesei*. Previously, 10-27% reduction of α-glucan, mannoprotein, β-glucan and chitin were observed in the *A. fumigatus ΔmsdS* mutant, while 25%, 33% and 55% in α-glucan, β-glucan and chitin respectively was observed when the temperature was increased (Li et al, 2008). On the other hand, the cell wall glycoprotein was reduced in Tr-MsdS strain (Fig.3C and Table S1). Previous investigation in *A. fumigatus* showed that unfolded protein response (UPR) is activated by reduced N-glycosylation and overexpressed cell wall protein and chitin (Li et al, 2011). In this study, UPR transcription activator factor *hac1*, was found to be reduced (Fig.4). This could provide some information that UPR is not induced in Tr-MsdS strain and there is no protein misfolding, which could have improved the growth and secretion in Tr-MsdS strain followed by enhanced α/β-glucan content. In addition, the enhanced expression of *sec61* and *rho3* which function to mediate protein translocation across ER and control the cell shape, respectively, has been clearly observed in Tr-MsdS strain. This provides support to the result that there is a change of the genes involved in secretory pathway of *T. reesei* and hence cell wall component has been altered when the *msdS* gene was introduced.

**Fig.4.**
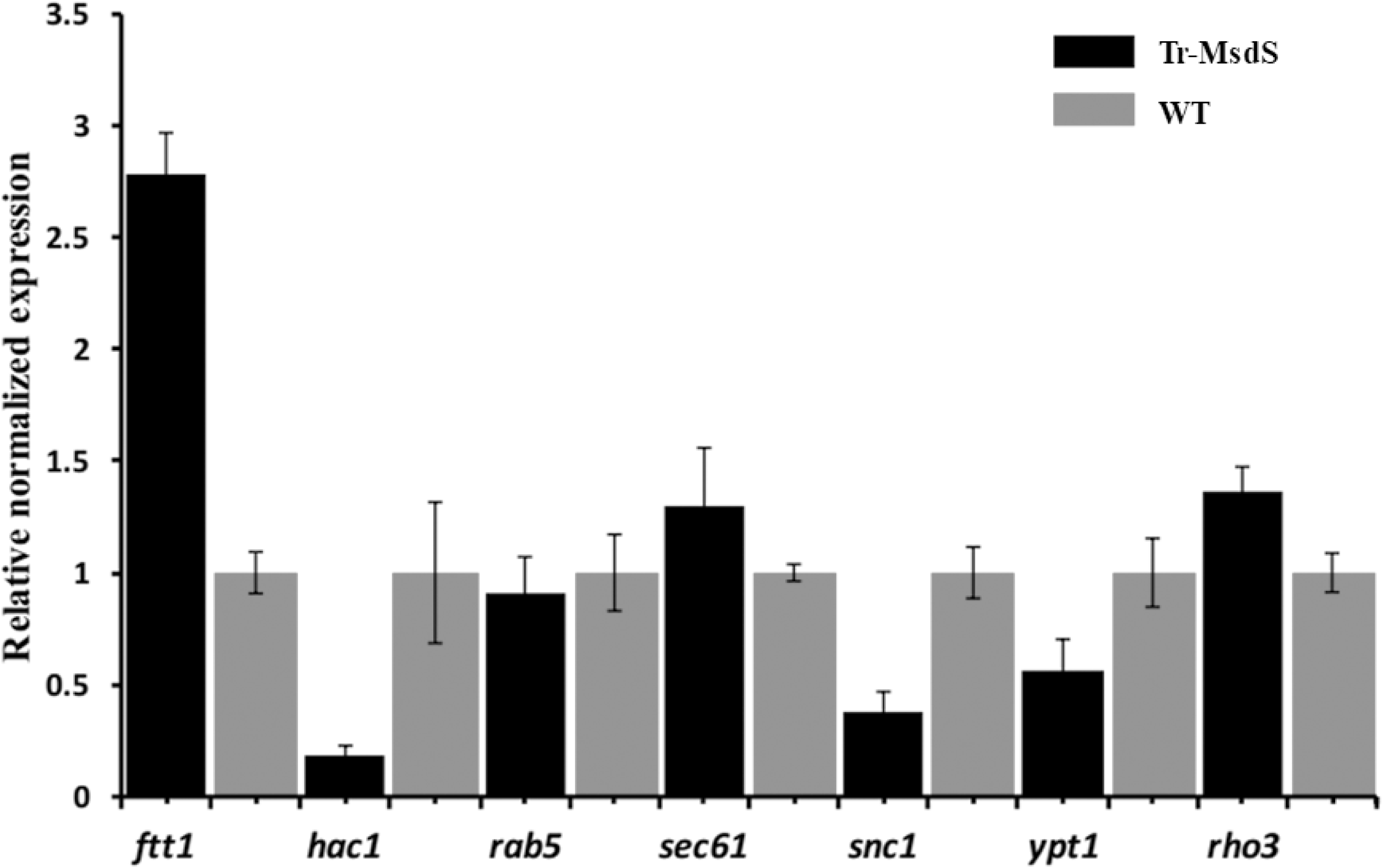
Activation of the genes involved in protein transport in *T. reesei*. 10^6^-10^7^ conidia were inoculated 100 ml of complete liquid medium and cultured at 32 °C. Mycelia were harvested and disrupted by grinding. Total RNA was extracted and quantification of mRNA levels were performed as described under method section. The 18s rRNA gene was used to standardize the mRNA levels of the target genes. Primers used in this assay are listed in Table S3. Each assay and each experiment were repeated 3 times. Results were presented as mean ± SD.

**Table 2.** Cell wall components of TrMsdS strain at 32°C and 37°C.

| Temp | Strain | Alkali soluble |  | Alkali insoluble |  |
| --- | --- | --- | --- | --- | --- |
|  |  | glycoprotein(μg) | α-glucan (μg) | chitin (μg) | β-glucan (μg) |
| 32°C | WT | 104.2±2.8 (100) | 215.2±1.1 (95.9) | 202.9±7.6 (100) | 268.2±1.7 (81.4) |
|  | Tr-MsdS | 81.6±0.3 (71.3) | 224.0±3.1 (100) | 175.1±2.6 (86.6) | 298.4±3.1 (100) |
| 37°C | WT | 185.4±14.8 (100) | 228.0±3.0 (94.4) | 282.4±3.9 (100) | 329.2±3.8 (78.2) |
|  | Tr-MsdS | 154.8±3.4 (61.3) | 241.4±1.9 (100) | 230.9±8.3 (89.7) | 363.5±1.7 (100) |

Cell wall components were isolated and determined as described by Yan et al. (55). Ten milligrams of lyophilized mycelia were used as independent samples for cell wall analysis with three different biological repeats. The values shown are microgram of cell wall component per 10 mg dry mycelia (±SD).

When the dormant conidia start to germinate in *A. fumigatus*, a series of nuclear division occur which is accompanied with an order of morphological events (switch from isotropic to polar growth) including appearance of first/second germ tubes and septation (Wang et al, 2015; Momany and Taylor, 2000). In the *A. fumigatus ΔmsdS* mutant random budding and septation at an early stage of germination were observed (Li et al, 2008). In *A. nidulans*, the temperature-sensitive mutant which is unable to switch from isotropic to polar growth was also reported although multiple points of polarity were established (Wang et al, 2015; Momany and Taylor, 2000). In some other studies, the abnormal morphology with balloon-like structures, swollen hyphae tips and altered cell cycle was described with a chitin synthase deficient mutant of *F. oxysporum* as well as *T. reesei* (Madrid et al, 2003).

In the present study, the parent and Tr-MsdS strain were grown in media containing glucose as the carbon source and observed for the germination of conidium. Fig.5 and Table 3 show that both parent and Tr-MsdS strain produce first germ tube at 6 h and initiation of the second germ tube at 7 h. In parent strain the germination showed a typical and regular branched apical growth usually at the early stage of germination, while less branched hyphae at the later stage of germination. However, in Tr-MsdS strain slightly swollen hyphal tips were observed together with multiple budding sites and random branching. In addition, during early stage, the first germ tube grew longer and then lateral germ tube appeared in Tr-MsdS strain. The spores of parent strain germinated in a typical polarized pattern in which the first and second germ tube occurred at 5 h and 6 h, respectively, and the first septation occurred after 9 h. Surprisingly, the occurrence of the first and second germ tube was rapid until 8 h in parent strain and then slowed down, whereas Tr-MsdS strain started to generate the second germ tube more rapidly after 8 h. The chitin accumulation was observed at the basal and lateral portion of hyphae. Very few septation occurred in both parent and Tr-MsdS strain. Previous studies have also reported that there is a proportional correlation of increased level of chitin with the level of glucan depicting that the cells will turned on the cell wall compensatory mechanism as a reaction under stress (Perlińska-Lenart et al, 2006). These results clearly demonstrated that insertion of the *msdS* gene led to more branched and random budding at later stage of germination.

**Fig.5.**
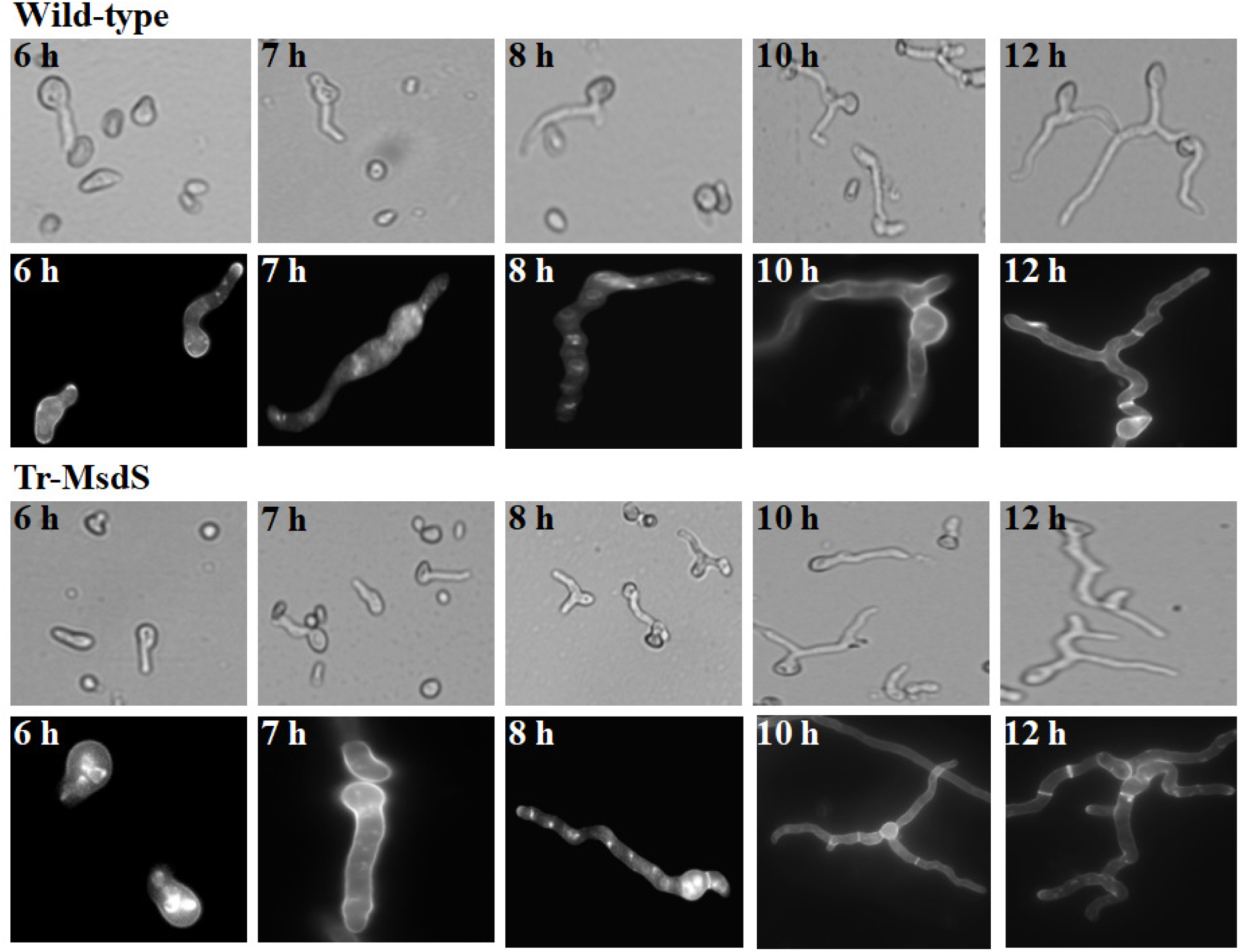
Germination of the wild-type and TrMsdS strain. Freshly harvested conidia were inoculated on petri dish containing 10 ml minimal media and glass coverslips and the adhering germinated conidia were observed under microscope (40X) at the specified times. The germlings of the wild-type (WT) and TrMsdS strain grown and fixed as described under method section. The nuclei and cell wall of the germlings were stained with DAPI and Calcofluor white respectively (100X).

**Table 3.** Statistics of germination of TrMsdS strain.

| Time<br>(h) | Number of germ tube (Tr-MsdS strain) |  |  |  | Number of germ tube (Wild-type strain) |  |  |  |
| --- | --- | --- | --- | --- | --- | --- | --- | --- |
|  | 0 | 1 | 2 | Hyper-<br>branching | 0 | 1 | 2 | Hyper-<br>branching |
| **5** | 156±9.5 | 3±1 | 0 | 0 | 149±15.7 | 3.33±1.5 | 0.3±0.6 | 0 |
| **6** | 100±10 | 6.7±1.2 | 0.7±0.6 | 0 | 87.3±16.3 | 12±6.3 | 1±1.4 | 0 |
| **7** | 78.3±4.7 | 23±3.6 | 1.7±1.5 | 0 | 88.7±4.2 | 27.3±2.1 | 2±1 | 0 |
| **8** | 65.3±3.5 | 35±2.7 | 9±2.7 | 0 | 64±3.6 | 33±4.4 | 4.3±1.5 | 0.3±0.6 |
| **9** | 39.3±2.5 | 44.7±4.5 | 19±2.7 | 2±1 | 51.7±3.1 | 39±4 | 8.7±1.5 | 1.3±0.6 |
| **10** | 10.7±2.5 | 46±10.8 | 30±3.6 | 3.3±0.6 | 12.3±1.5 | 48±2.7 | 16±2.7 | 1.7±0.6 |
| **12** | 7±2 | 21±7.6 | 43.7±2.5 | 7±2 | 6.3±3.8 | 28.67±2.1 | 28.33±4.4 | 5.7±3.1 |

Freshly harvested conidia were poured into a Petri dish containing glass coverslips and incubated in 10 ml of liquid medium. The adhered germlings on the coverslips were counted for number of germ tubes appear during different stages of germination under microscope. For each independent experiment approximately 100 conidia were counted and three independent experiments were carried out.

### Glycosylation and protein expression in Tr-MsdS strain

To compare the N-glycosylation between parent and mutant strain, the N-glycans were released from membrane and secreted proteins with PNGase F and analyzed by MALDI-TOF. As shown in Fig.6, signals of Man_5-8_GlcNAc_2_ were detected in parent strain, among them Man_8_GlcNAc_2_ was the major glycoform. Although signals corresponding to Man_5-9_GlcNAc_2_ were detected in Tr-MsdS strain, the glycoform Man_8_GlcNAc_2_ was greatly reduced, suggesting an action of MsdS on N-glycans.

**Fig.6.**
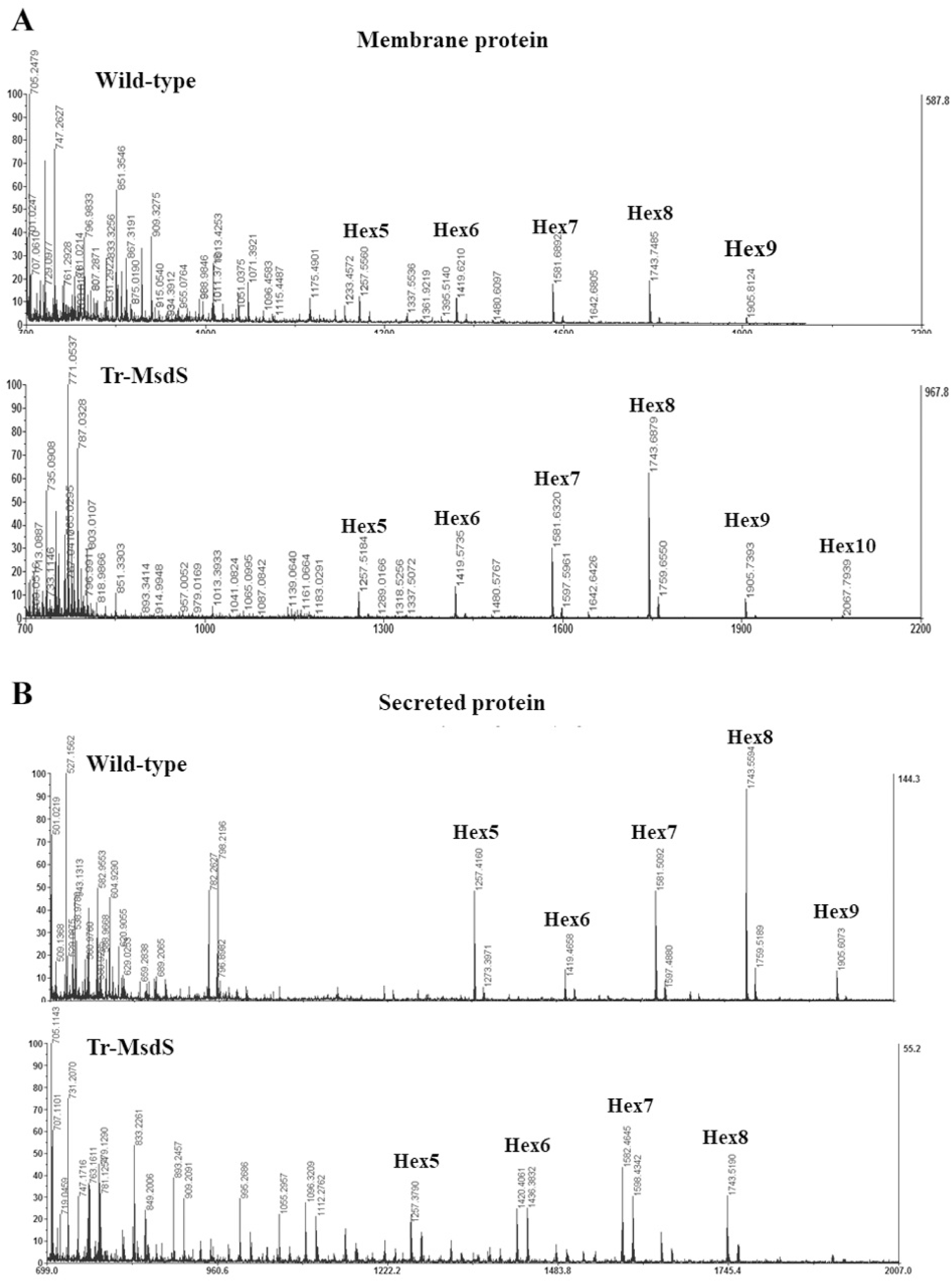
Detection of N-glycans on membrane and secreted proteins. N-glycans from membrane proteins (A) and secreted proteins (B) were released by PNGase F and subjected to Mass spectrometry analysis.

As shown in Fig.2B, the intracellular, cell wall, membrane and secreted proteins of both strains showed no difference. However, the MS/MS analysis of secreted protein identified some important proteins involved in secretory and regulatory pathway of *T. reesei*, such as 14-3-3-like protein (gi|12054274), HSP70 (gi|30961863) and small GTPase of the Rab/Ypt family (gi|340519278) (Table S2). Saloheimo and Pakula have clearly explained about the secretory pathway in filamentous fungi including *T. reesei* (Saloheimo and Pakula, 2012). Based on the *T. ressei* protein secretory pathway (Fig.7), the results were further analyzed by transcriptional expression (Fig.4). Interestingly, it was observed that three genes encoding Sec61 (major component of protein translocation complex in ER membrane), Ftt1 (a 14-3-3 type involved in last step of secretory pathway) and Rho3 (Ras-type GTPase involved in cell polarity and vesical fusion with plasma membrane) were found to be expressed higher in Tr-MsdS strain as compared with its parent strain. Whereas expressions of the genes encoding Hac1, Rab5, Snc1 and Ypt1 were reduced in Tr-MsdS strain. These results suggested that expression of MsdS affected the protein transportation and secretion in *T. reesei*.

**Fig.7.**
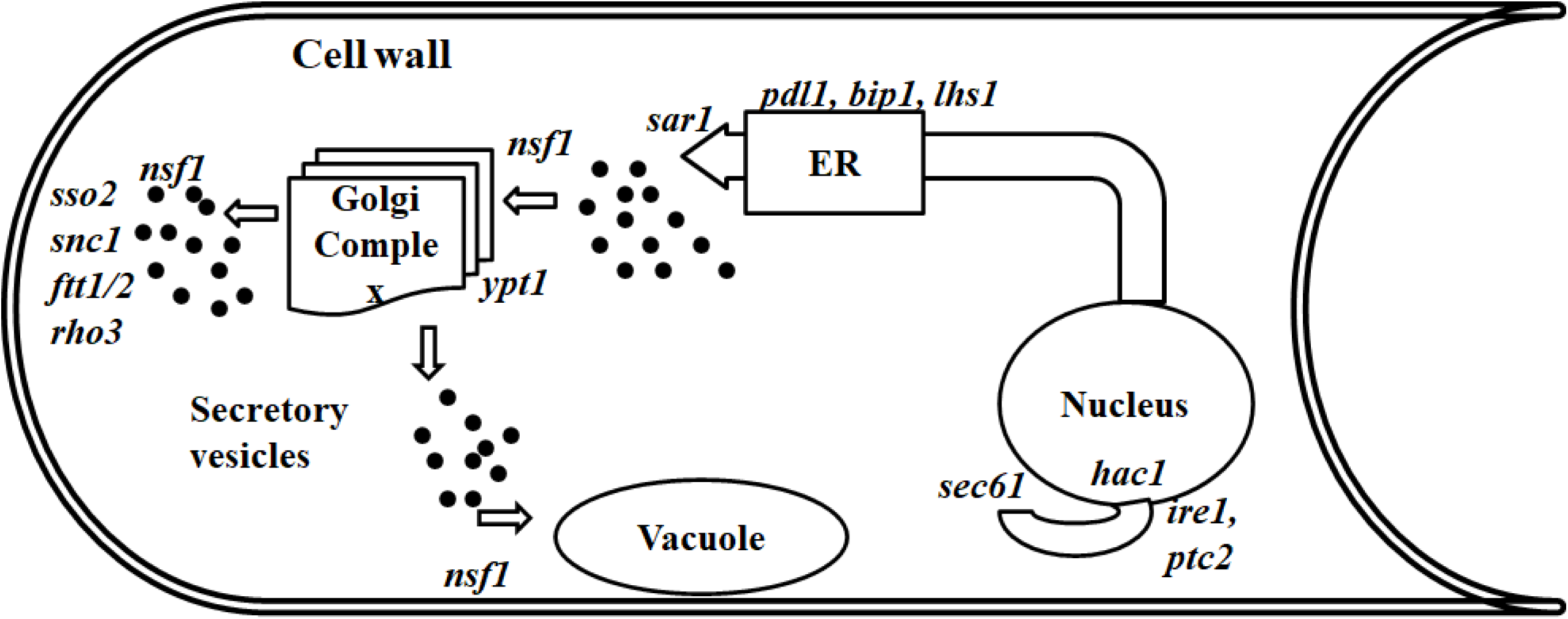
Schematic of the secretory pathway in *T. reesei*. The pathway components constitute gene characterized in *T. reesei*. *hac1*, transcription factor of the UPR; *ire1*, sensor of UPR; *ptc2*, phosphatase acting as a negative regulator of UPR; *sec61*, major component of the protein translocation complex in ER membrane; *pdi1*, protein disulphide isomerase; *bip1* and *lhs1*, HSP70 family chaperones involved in protein folding in ER; *sar1*, small GTPase involved in vesicle budding from the ER; *ypt1*, Ras-type small GTPase involved in vesicle fusion into Golgi; *nsf1*, general fusion factor involved in multiple vesicle fusion steps; *snc1*, v-SNARE protein involved in vesicle fusion to the plasma membrane; *sso1/2*, t-SNARE proteins involved in vesicle fusion to the plasma membrane; *ftt1/2*, 14-3-3 type proteins involved in the last step of the secretory pathway; and *rho3*, Ras-type small GTPase involved in cell polarity and vesicle fusion with plasma membrane.

### Cellulase and β-mannanase in Tr-MsdS strain

In the present study, cellulase and β-mannanase activity were analyzed using carboxymethyl cellulose and locust bean gum as a substrate, respectively. Both activity and hydrolysis yield were found to be higher in Tr-MsdS strain as compare with its parent strain. As shown in Fig.8, glucose yield was 17.5%, 12.6% and 9.9% higher in Tr-MsdS strain at 30°C, 40°C and 50°C, respectively. Also, mannose yields in Tr-MsdS strain were 27.1%, 25.7% and 32.2% higher than that in the parent strain at 30°C, 40°C and 50°C, respectively. These observations suggest that introduction of MsdS into *T. ressei* not only affects the glycosylation and cell wall synthesis, but also improves the ligno-cellulose degradation. Further, the yield was found to be increased at temperature up to 50°C (industrially applicable temperature). This could also somehow attribute to be applied in industrial purpose to degrade complex substrate after optimizing proper condition for efficient degradation. The Tr-MsdS in the present study somehow acted as a fungus constructed with additional mannan degrading enzyme providing improved hydrolysis of locust bean gum or cellulose to some extent.

**Fig.8.**
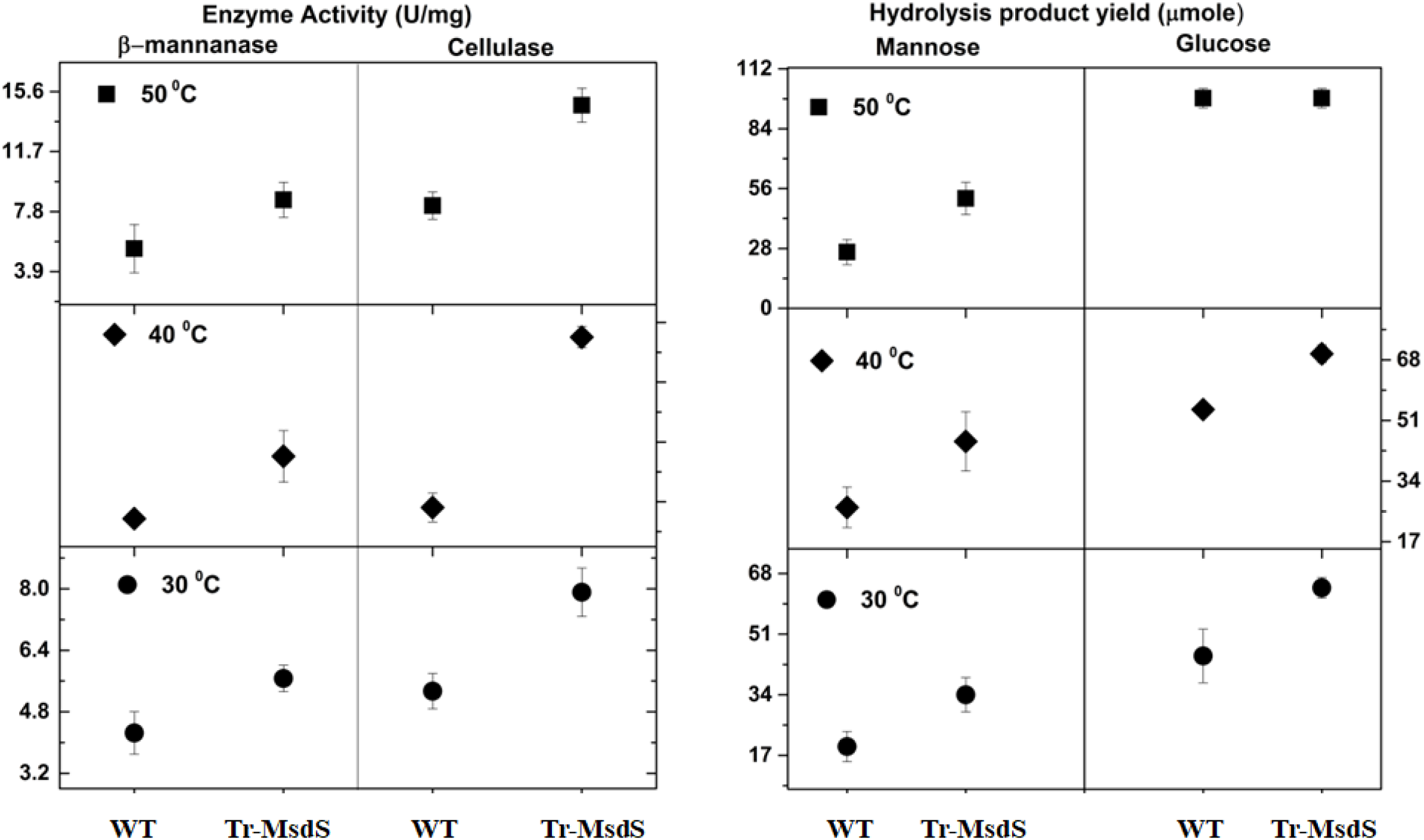
Hydrolysis yield and enzyme activity of secreted protein produced by Tr-MsdS strain. The secreted proteins obtained from both wild-type and Tr-MsdS strains were mixed with locust bean gum and CMC-cellulose. After reaction the release of sugar (mannose or glucose) (A) and enzyme activity (β-mannanase or cellulase, respectively) (B) were measured. Results were presented as mean ± SD.

## Discussion

The primary steps of protein N-glycosylation is common among fungi and mammals including the site-specific transfer of Glc_3_Man_9_GlcNAc_2_ from ER lumen to the de novo synthesized protein followed by subsequent trimming by glucosidases I and II and a specific ER-residing α−1,2-mannosidase to form Man_8_GlcNAc_2_ structures (isomer Man8B), which later export the predominant Man8-isomer to the Golgi (Choi et al, 2003). In human Golgi α-1,2-mannosidase (IA-IC) removes Man to yield the Man_5_GlcNAc_2_ structure (the precursor for complex N-glycans) (Wildt and Gerngross, 2005; Gleeson, 1998; Jacobs et al, 2009), whereas in *S. cerevisiae* (Wildt and Gerngross, 2005), the N-glycan processing, involves the addition of numerous mannose sugars throughout the entire Golgi, often leading to hyper-mannosylated N-glycan structures. Previous studies have explained the consequences of Golgi α-mannosidase I activity vary in different species. The deficiency of α-mannosidase results in the lethal disease called mannosidosis in human (Khan and Ranganathan, 2009) and cattle (Phillips et al, 1974), and outer chain synthesis in *S. cerevisiae* (Puccia et al, 1993).

In *T. reesei*, the most common N-glycan structure is found to be mono-glucosylated high-mannose glycan GlcMan_7-8_GlcNAc_2_, which is similar to a transient structure of GlcMan_9_GlcNAc_2_ found in the ER (Stals et al, 2004). However, a number of other N-glycan structures also have been reported in different strains. For example, the N-glycan of Cel7A in *T. reesei* strain QM9414 and ALKO2877 is single GlcNAc residues in three (Asn45, Asn270, and Asn384) out of four potential sites characterized in the catalytic domain (Harrison et al, 1998), while Garcia et al. also identified heterogeneous N-glycans on endoglucanase I (Cel7B) from strain QM9414 (García et al, 2001). In hyper-producing strain Rut-C30, more complex N-glycan structures, including single GlcNAc and Hex_8-9_GlcNAc_2_ structures, are found on cellulasese Cel7B, Cel6A and Cel5A (Maras et al, 1997; Hui et al, 2002). These observations suggest that the N-glycan structures on cellulases produced by different mutant strains are different. Although *T. reesei α*-1,2-mannosidase can sequentially cleave all α-1,2-linked mannose sugars from a Man_9_GlcNAc_2_ oligosaccharide giving broad substrate specificity that sterically allowed different oligosaccharide conformations (Van Petegem et al, 2001), it is also possible that some enzymes in glycosylation pathway or some glycosylation sties on cellulases were mutated as these mutants were screened for cellulases production or high-yield of protein expression. Somehow, it is no doubt that N-glycosylation is involved in protein secretion.

In contrast to the Hex_8-9_GlcNAc_2_ structures in *T. reesei*, the major glycoform N-glcan in *A. fumigatus* is Man_6_GlcNAc_2_ (Li et al, 2008). In *A. fumigatus*, deletion of the Golgi α-mannosidase gene *msdS* causes defective N-glycan processing and gives rise to Man_8_GlcNAc_2_ glycoform, which is similar with that of the wild-type *T. reesei*. Interestingly, the conversion of glycoform from Man_6_GlcNAc_2_ to Man_8_GlcNAc_2_ leads phenotypes such as defective cell wall, reduced conidiation and abnormal polarity in *A. fumigatus* (Li et al, 2008), suggesting a species-specific N-glycan structure in different filamentous fungi. To verify this hypothesis, in this study we introduced the *A. fumigatus msdS* gene into *T. reesei*.

The *gpdA* gene encodes glyceraldehyde-3-phosphate dehydrogenase (GPD) and is a key enzyme in glycolysis and glucogenesis which constitutes up to 5% of the soluble cellular protein in *Saccharomyces cerevisiae* (Krebs et al, 1953; Redkar et al, 1998) and *A. nidulans* (Redkar et al, 1998). Several copies of GPD-encoding genes have been accounted in higher eukaryotes (Redkar et al, 1998; Fort et al, 1985), but only a single GPD-encoding gene has been reported in *A. nidulans* (Punt et al, 1988). The *gpdA*-promoter-controlled exocellular production of glucose oxidase by recombinant *A. niger* NRRL-3 during growth on glucose and non-glucose carbon sources was investigated for better carbon substrate identification (El-Enshasy et al, 2001). As several studies have successfully reported the gene manipulation in *T. reesei* under the *A. nidulans gpdA* gene promoter and *trpC* (indole-3-glycerol phosphate synthase) terminator (Zakrzewska et al, 2003), we therefore generate a vector carrying *gpdA* promoter and *msdS* gene. The resulting vector was co-transformed into *T. reesei* with plasmid TAkulox with *pyr4* gene as a selective marker.

Analysis of Tr-MsdS strain revealed that, although its cell wall integrity was not affected, expression of MsdS in *T. reesei* led to morphological changes including increased cell wall thickness, abnormal polarity and ballooned hyphal tips. Further analysis demonstrated that the genes encoding proteins involved in protein translation, folding and vesicle transportation were induced, such as Sec61, HSP70, Rab/Ypt family GTPase, Rho3 and Ftt1, indicating that alteration of the N-glycan processing by MsdS affects the protein transportation and secretion in *T. reesei*. These results demonstrate that N-glycan processing is required for protein secretion and species-specific in filamentous fungi.

Filamentous fungi are thought as the most efficient lignocelluloses degraders, especially *T. reesei* (Zhang et al, 2015; Wang et al, 2014), however, there is still an insufficient hydrolysis of lignocellulose due to the moderately low levels of mannan-degrading activity (Inoue et al, 2015). To compensate for the lower mannan-degrading activity, exogenous mannosidases have been added in the cellulase system. For example, in *Talaromyces cellulolyticus* the approximately 80% mannose yield was increased with high β-mannanase and β-mannosidase activities when supplemented with *A. niger* derived commercial enzyme (Inoue et al, 2015). It has been shown that the crystal of mannan can be degraded when endo-α-1,4-mannanase from *T. reesei*, endo-α-1,4-mannanase from *A. niger* and exo-α-1,4-mannosidase from *A. niger* are used (Hagglund et al, 2001). To evaluate the effects of N-glycan processing on protein secretion, in this study we further analyzed cellulase and β-mannanase activities in Tr-MsdS strain. Our results showed that both cellulase-degradation and mannan-degradation activities were increased by 10-32% (Fig.8) in Tr-MsdS strain, confirming that introduction of MsdS into *T. ressei* affects protein secretion and improves the ligno-cellulose degradation.

Overall, although growth rate and cell wall integrity were not altered, phenotypes such as increased cell wall components, increased conidiation, abnormal polarity and enhanced cellulase activity were documented in strain Tr-MsdS. In conclusion, our results confirm that N-glycan processing between *A. fumigatus* and *T. reesei* is species-specific and required for protein secretion. In addition, our results also provide a new strategy to improve cellulases production by interfering the N-glycan processing in *T. reesei*.

## Materials and methods

### Strains and growth conditions

*T. reesei* TU-6 (uridine auxotrophs, American Type Culture Collection, ATCCMYA-256), obtained from Prof. Zhiyang Dong, Institute of Microbiology, Chinese Academy of Sciences), was used as the recipient of the α-mannosidase gene from *A. fumigatus*. TU-6 was cultivated in minimal medium (MM Medium) per liter containing 2% glucose, 0.5% (NH_4_)_2_SO_4_, and 1.5% KH_2_PO_4_. Microelements (0.005% FeSO_4·_7H_2_O, 0. 0016% MnSO_4·_7H_2_O, 0.0014% ZnSO_4·_7H_2_O, 0.0037% CoCl_2_, 0.6% CaCl_2_ and 0. 6% MgSO_4·_7H_2_O) and 7 mmol uridine was also added in the medium to culture TU-6 at 28°C on a rotary shaker (200 rpm). Conidia were prepared by cultivating *T. reesei* TU-6 strains on solid PDA medium with uridine for 5-7 days at 28°C. The spores were collected, washed thrice with and resuspended in sterilized ddH_2_O (added with 40% glycerol solution (1:1) for long term storage at −70°C. The spore concentration was confirmed using haemocytometer counting and viable counting.

### Expression of the *A. fumigatus msdS* in *T. reesei*

*Escherichia coli* DH5α was used for plasmid propagation in order to insert the α-mannosidase *msdS* gene from *A. fumigatus* in TU-6. pCG (5.5 kb vector plasmid containing *chiB* promoter and GFP) vector was reconstructed into pGM vector by using GPDA as a promoter with oligonucleotides; *gpda*-F (CAATTCCCTTGTATCTCTACACACAG) with *EcoR*I restriction site and *gpda*-R (GGTGA TGTCTGCTCAAGCG) with *KpnI* restriction site and *msdS* as insert gene with oligonucleotides; *MsdS*-F (ATGCATTTACCCTCTTTGTCCGTG) and *MsdS*-R (TCACGTATGATGAATTCGGACAGGGTG) as forward and reverse primers at *Kpn*I and *BamH*I restriction site respectively. Both *gpda* and *msdS* were cloned into pGEM-T Easy (Promega, USA) and confirmed by sequencing. And then inserted into the pCG vector at specific site and confirmed by PCR and sequencing. For the transformation of plasmid into *T. reesei* TU-6 protoplast, a co-transformation with TAkulox *pyr4* group containing plasmid was used as a selectable marker. The transformants were regenerated in MM media containing sorbitol (1 mol/L D-sorbitol). The transformants were screened in MM media containing 1.5% agar and confirmed by PCR and sequencing.

### Protein extraction

The extraction of proteins from different fractions of cells was done as described by Wang *et al*. (Wang et al, 2015) with some modifications. The extracellular proteins were precipitated with freshly prepared 2% (w/v) sodium deoxycholate to the culture supernatant (1:100) for 30 min on ice followed by addition of 100% trichloroacetic acid (1:10) for 30 min on ice, and collected by centrifugation (12,000 × g at 4°C for 10 min). The precipitate was washed 3 times with ice-cold acetone (once with 100% [v/v] and twice with 90% [v/v]) and dried by exposure to air. For cytosolic and membrane proteins extraction, the mycelia obtained from culture was ground with buffer I (200 mM Tris–HCl, 50 mM EDTA, protease inhibitor cocktail) at 4°C for 2 h, centrifuged at 4,000g for 10 min and the supernatant obtained (containing cytosolic and membrane proteins) was ultra-centrifuged 50,000g at 4°C for 1 h. After centrifugation, the supernatant containing cytosolic proteins fraction and the precipitate containing membrane protein fraction were collected separately. The mycelia obtained from the culture was dried, weighted, and resuspended in 25 µl of Tris buffer (0.05 M Tris–HCl, pH 7.8) per mg of dried mycelia to extract cell wall protein. After centrifugation the pellets obtained were boiled three times in 25 µl of sodium dodecyl sulfate extraction buffer (50 mM Tris–HCl, 2% SDS, 20 mM Na-EDTA, and 40 mM β-mercaptoethanol) per mg of dried mycelium followed by centrifugation (12,000 r.p.m) at 4°C for 10 min. The pellet was washed with distilled water three times, dried, and treated with 10 µl of hydrofluoride (HF)-pyridine per gram mycelium and put on ice for 3 h. The supernatant was separately collected and added with 9 volumes of 100% methanol buffer (100% methanol, 50 mM Tris–HCl, pH 7.8) at 0°C for 2 h. Finally, the cell wall proteins were collected by centrifugation (12,000 r.p.m) at 4°C for 10 min after washing with 90% methanol buffer (90% methanol, 50 mM Tris–HCl, pH 7.8) three times. The concentration of each fraction of proteins was determined by SDS-PAGE and quantitatively by Bradford protein assay (BIO-RAD, USA).

### SDS-PAGE and in-gel digestion

The proteins were separated by 10-12% SDS-PAGE, stained with R-250, and then cut out from gel. In-gel digestion of secreted protein in SDS-PAGE was performed according to Liu *et al.* with a slight modification (Liu et al, 2013). Briefly, each sliced band was de-stained by using 50% (v/v) acetonitrile in 40 mM NH_4_HCO_3_, dehydrated using 100% acetonitrile and dried using SpeedVac. Proteins were then reduced with 10 mM DTT/40 mM NH_4_HCO_3_ (Sigma-Aldrich Co.) at 56°C for 20 min, alkylated with 55 mM iodoacetamide/40 mM NH_4_HCO_3_ in the dark for 25 min at room temperature followed by subsequent wash and dry. Peptides were then produced by trypsin digestion (50 ng trypsin, Promega Sequence Grade, Modified) (Liu et al, 2013). For MALDI-MS analysis, 0.4 μl aliquot of the reconstituted tryptic peptide mixture in 0.1% TFA was mixed with 0.4 μl of CHCA matrix solution (5 mg/ml CHCA in 50% ACN/0.1% TFA) and spotted onto a freshly cleaned target plate. After air drying, the crystallized spots were analyzed on the MALDI-TOF/TOF 5800 (AB SCIEX, Framingham, MA).

### Analysis of N-glycan

N-glycans were released from membrane and secreted proteins of *T. reesei* strain by peptide N-glycosidase F (PNGase F, NEB, P0704) (Wang et al, 2015). The enzyme reaction includes the process of denaturation of proteins at 95°C for 5 min followed by addition 10% Nonidet P40 (NP40) treating with PNGase F. After digestion, the sample was centrifuged and the supernatant was subjected to C8 column, washed with 100% acetonitrile (ACN) and equilibrated with 0.1% trifluoroacetic acid (TFA) to separate N-glycans from proteins. The released N-glycans were collected and applied to a graphite column, washed with 0.1% TFA to remove salts and then eluted with elution buffer (60%ACN, 0.1% TFA) to collect N-glycans. The structure of released and purified N-glycans was analyzed by MALDI-TOF-MS.

### Phenotypic analysis

The growth kinetics was determined by measuring colony diameter. Same amount of spore was inoculated on media plate and the colony growth was monitored by measuring the diameter of each colony at different time intervals (Wang et al, 2015). Similarly, the sensitivity of the mutant to antifungal reagents was also analyzed (Li et al, 2008; Wang et al, 2015). The conidiophores were spotted on solid PDA medium with uridine (MMU) plates with same concentration in the presence of 100 µg/ml calcofluor white, 150 µg Congo red or 40 µg SDS and incubated at 28°C, 32°C and 37°C and the plates were analyzed for colony growth, measured for colony diameter and photographed if required.

### Electron Microscopy

Scanning and transmission electron microscopy analyses were performed as described by Li et al. (Li et al, 2008). Culture samples were fixed with 2.5% glutaraldehyde in phosphate buffer pH 7.2 and then examined with FESEM HITACHI SU8010 scanning electron microscope. While for TEM, culture sample were fixed with 2.5% glutaraldehyde in 0.1 M phosphate at room temperature or 4°C for 2 h or overnight, washed three times with 0.1 M phosphate buffer, post-fixed in 1% osmium tetroxide, incubated at room temperature for 2-4 h in 0.1 M phosphate. The cells were washed with 0.1 M phosphate buffer 2-3 times and then dehydration was done by treating with each of 30, 50, 70, 85, 95, and 100% methanol for 15-20 min each. Infiltration was done by using LR white resin: ethanol (1:1) for 4 h followed by rinsing with LR white resin (100%) overnight once and rinsed for 2 h again. Cells were embedded in mold at 55°C for 24 h followed by sectioning (Leica uc7) and staining (uranyl acetate for 25 min and lead citrate for 5 min) and the sections were examined with JEM-1400.

### Germination

10^6^ freshly harvested conidia were inoculated in 10 ml of minimal liquid medium in a Petri dish containing 5-6 glass coverslips at 32°C. The coverslips with adhering germinated conidia were taken out and counted for the number of germ tubes germinated at the specified times counted under differential interference contrast microscopy (Li et al, 2008). For the observation of nuclei and septum the conidia adhered on the glass coverslip were fixed in fixative solution (4% formaldehyde, 50 mM phosphate buffer, pH 7.0, and 0.3% Triton X-100). After 30 min of fixation, coverslips were washed with phosphate-buffered saline (PBS), incubated with 1 µg/ml 4,6-diamidino-2-phenylindole (DAPI) (Sigma) for 20 min and washed with PBS. Then the germinated conidia on glass coverslips was dyed with a 5 µg/ml solution of fluorescent brightener 28 (Sigma) for 5 min and washed with PBS for 5 times (Wang et al, 2015; Fang et al, 2013). The conidia were then observed and photographed using a fluorescence microscope (Axiovert 200 M, Carl Zeiss).

### Quantitative Real Time PCR

One hundred milliliters of complete liquid medium were inoculated with 10^6^-10^7^ conidia and the harvested mycelia were disrupted by grinding. Total RNA was extracted using TRIzol reagent (Invitrogen/Life Technologies, Carlsbad, CA, US) (Wang et al, 2015; de Souza et al, 2011). The cDNA synthesis was carried out with 5 µg RNA using the RevertAid™ First Strand cDNA Synthesis Kit (Fermentas). The forward and reverse primers used for cDNA synthesis was *MsdS*-FGCTTGCTTTGATGGAGGAAG and *MsdS*-R TGACGCGGTACGCATAGTAG, respectively. The quantitative PCR reaction was done with SYBR®PremixExTaq™ (Takara) with the thermal cycling condition of 95°C for 30 s, followed by 40 cycles of 95°C for 5 s and 60°C for 30 s (Bio-Rad CFX manager 3.1). Quantification of mRNA levels of different genes was performed using the 2^−ΔΔct^ method. The 18s rRNA gene was used to standardize the mRNA levels of the target genes. Each assay and each experiment were repeated 3 times. To avoid or detect any possible contamination or carryover, appropriate negative controls containing no template were also subjected to the same procedure. Primers used in this study are listed in Table S3.

### Enzyme activity assays and hydrolysis yield

The secreted proteins from different strains were analyzed for cellulase and β-mannanase activity as well as reducing sugar yield at three different condition of temperature (30°C, 40°C and 50°C) using carboxymethyl cellulose or locust bean gum as a substrate (Sigma-Aldrich Co.). The activity assay was determined by using 3,5-dinitrosalicylic acid (DNS) method (Dashtban et al, 2010; Dale et al, 1996). The reaction was started by mixing 5 mg/ml carboxymethyl cellulose or 5 mg/ml locust bean gum in 0.1 M citrate-phosphate buffer with 0.1 ml secretory protein sample and incubated at three different temperatures. One unit of cellulase or endo-1,4-β-mannosidase activity is defined as amount of enzyme releasing 1 µmole of glucose or 1 µmole mannose equivalent per minute and the hydrolysis yield (µmole) of reducing sugar was measured by measuring the amount of reducing sugar produced from the reaction. Triplicates of each sample were analyzed and mean values were calculated.

## Acknowledgements

This work is sponsored by the Natural Science Foundation of China (31630016 and 31320103901) and partially by Bagui Scholar Program Fund (2016A24) of Giuangxi Zhuang Autonomous Region to CJ.

